# An efficient not-only-linear correlation coefficient based on machine learning

**DOI:** 10.1101/2022.06.15.496326

**Authors:** Milton Pividori, Marylyn D. Ritchie, Diego H. Milone, Casey S. Greene

## Abstract

Correlation coefficients are widely used to identify patterns in data that may be of particular interest. In transcriptomics, genes with correlated expression often share functions or are part of disease-relevant biological processes. Here we introduce the Clustermatch Correlation Coefficient (CCC), an efficient, easy-to-use and not-only-linear coefficient based on machine learning models. CCC reveals biologically meaningful linear and nonlinear patterns missed by standard, linear-only correlation coefficients. CCC captures general patterns in data by comparing clustering solutions while being much faster than state-of-the-art coefficients such as the Maximal Information Coefficient. When applied to human gene expression data, CCC identifies robust linear relationships while detecting nonlinear patterns associated, for example, with sex differences that are not captured by linear-only coefficients. Gene pairs highly ranked by CCC were enriched for interactions in integrated networks built from protein-protein interaction, transcription factor regulation, and chemical and genetic perturbations, suggesting that CCC could detect functional relationships that linear-only methods missed. CCC is a highly-efficient, next-generation not-only-linear correlation coefficient that can readily be applied to genome-scale data and other domains across different data types.

## Introduction

New technologies have vastly improved data collection, generating a deluge of information across different disciplines. This large amount of data provides new opportunities to address unanswered scientific questions, provided we have efficient tools capable of identifying multiple types of underlying patterns. Correlation analysis is an essential statistical technique for discovering relationships between variables [1]. Correlation coefficients are often used in exploratory data mining techniques, such as clustering or community detection algorithms, to compute a similarity value between a pair of objects of interest such as genes [2] or disease-relevant lifestyle factors [3]. Correlation methods are also used in supervised tasks, for example, for feature selection to improve prediction accuracy [4,5]. The Pearson correlation coefficient is ubiquitously deployed across application domains and diverse scientific areas. Thus, even minor and significant improvements in these techniques could have enormous consequences in industry and research.

In transcriptomics, many analyses start with estimating the correlation between genes. More sophisticated approaches built on correlation analysis can suggest gene function [6], aid in discovering common and cell lineage-specific regulatory networks [7], and capture important interactions in a living organism that can uncover molecular mechanisms in other species [8,9]. The analysis of large RNA-seq datasets [10,11] can also reveal complex transcriptional mechanisms underlying human diseases [2,12,13,14,15]. Since the introduction of the omnigenic model of complex traits [16,17], gene-gene relationships are playing an increasingly important role in genetic studies of human diseases [18,19,20,21], even in specific fields such as polygenic risk scores [22]. In this context, recent approaches combine disease-associated genes from genome-wide association studies (GWAS) with gene co-expression networks to prioritize “core” genes directly affecting diseases [19,20,23]. These core genes are not captured by standard statistical methods but are believed to be part of highly-interconnected, disease-relevant regulatory networks. Therefore, advanced correlation coefficients could immediately find wide applications across many areas of biology, including the prioritization of candidate drug targets in the precision medicine field.

The Pearson and Spearman correlation coefficients are widely used because they reveal intuitive relationships and can be computed quickly. However, they are designed to capture linear or monotonic patterns (referred to as linear-only) and may miss complex yet critical relationships. Novel coefficients have been proposed as metrics that capture nonlinear patterns such as the Maximal Information Coefficient (MIC) [24] and the Distance Correlation (DC) [25]. MIC, in particular, is one of the most commonly used statistics to capture more complex relationships, with successful applications across several domains [4,26,27]. However, the computational complexity makes them impractical for even moderately sized datasets [26,28]. Recent implementations of MIC, for example, take several seconds to compute on a single variable pair across a few thousand objects or conditions [26]. We previously developed a clustering method for highly diverse datasets that significantly outperformed approaches based on Pearson, Spearman, DC and MIC in detecting clusters of simulated linear and nonlinear relationships with varying noise levels [29]. Here we introduce the Clustermatch Correlation Coefficient (CCC), an efficient not-only-linear coefficient that works across quantitative and qualitative variables. CCC has a single parameter that limits the maximum complexity of relationships found (from linear to more general patterns) and computation time. CCC provides a high level of flexibility to detect specific types of patterns that are more important for the user, while providing safe defaults to capture general relationships. We also provide an efficient CCC implementation that is highly parallelizable, allowing to speed up computation across variable pairs with millions of objects or conditions. To assess its performance, we applied our method to gene expression data from the Genotype-Tissue Expression v8 (GTEx) project across different tissues [30]. CCC captured both strong linear relationships and novel nonlinear patterns, which were entirely missed by standard coefficients. For example, some of these nonlinear patterns were associated with sex differences in gene expression, suggesting that CCC can capture strong relationships present only in a subset of samples. We also found that the CCC behaves similarly to MIC in several cases, although it is much faster to compute. Gene pairs detected in expression data by CCC had higher interaction probabilities in tissue-specific gene networks from the Genome-wide Analysis of gene Networks in Tissues (GIANT) [31]. Furthermore, its ability to efficiently handle diverse data types (including numerical and categorical features) reduces preprocessing steps and makes it appealing to analyze large and heterogeneous repositories.

## Results

### A robust and efficient not-only-linear dependence coefficient

The CCC provides a similarity measure between any pair of variables, either with numerical or categorical values. The method assumes that if there is a relationship between two variables/features describing *n* data points/objects, then the **cluster**ings of those objects using each variable should **match**. In the case of numerical values, CCC uses quantiles to efficiently separate data points into different clusters (e.g., the median separates numerical data into two clusters). Once all clusterings are generated according to each variable, we define the CCC as the maximum adjusted Rand index (ARI) [32] between them, ranging between 0 and 1. Details of the CCC algorithm can be found in Methods.

We examined how the Pearson (*p*), Spearman (*s*) and CCC (*c*) correlation coefficients behaved on different simulated data patterns. In the first row of Figure 1, we examine the classic Anscombe’s quartet [33], which comprises four synthetic datasets with different patterns but the same data statistics (mean, standard deviation and Pearson’s correlation). This kind of simulated data, recently revisited with the “Datasaurus” [34,35,36], is used as a reminder of the importance of going beyond simple statistics, where either undesirable patterns (such as outliers) or desirable ones (such as biologically meaningful nonlinear relationships) can be masked by summary statistics alone.

**Figure 1:**
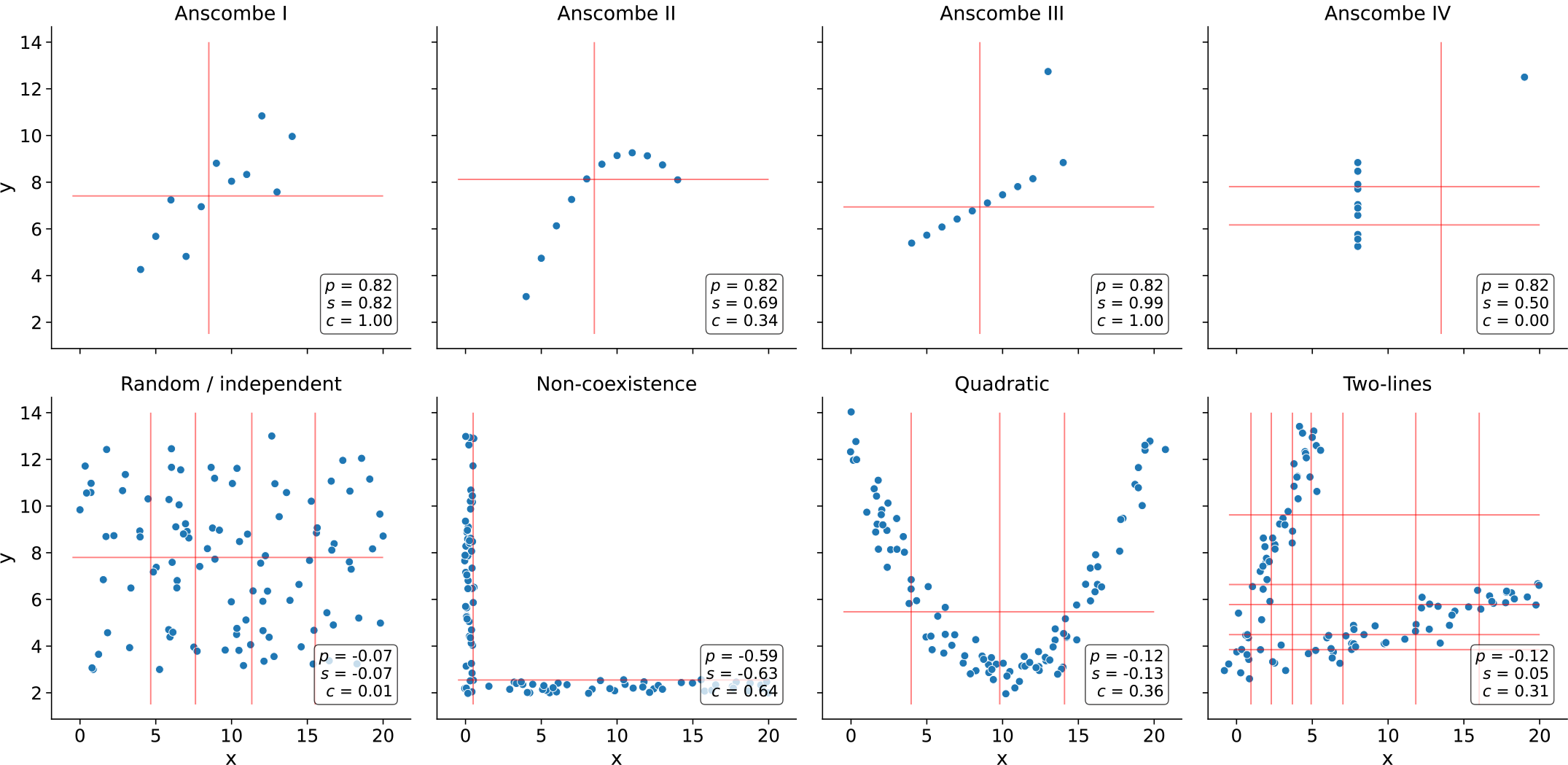
Different types of relationships in data. Each panel contains a set of simulated data points described by two generic variables: *x* and *y*. The first row shows Anscombe’s quartet with four different datasets (from Anscombe I to IV) and 11 data points each. The second row contains a set of general patterns with 100 data points each. Each panel shows the correlation value using Pearson (*p*), Spearman (*s*) and CCC (*c*). Vertical and horizontal red lines show how CCC clustered data points using *x* and *y*.

Anscombe I contains a noisy but clear linear pattern, similar to Anscombe III where the linearity is perfect besides one outlier. In these two examples, CCC separates data points using two clusters (one red line for each variable *x* and *y*), yielding 1.0 and thus indicating a strong relationship. Anscombe II seems to follow a partially quadratic relationship interpreted as linear by Pearson and Spearman. In contrast, for this potentially undersampled quadratic pattern, CCC yields a lower yet non-zero value of 0.34, reflecting a more complex relationship than a linear pattern. Anscombe IV shows a vertical line of data points where *x* values are almost constant except for one outlier. This outlier does not influence CCC as it does for Pearson or Spearman. Thus *c* = 0.00 (the minimum value) correctly indicates no association for this variable pair because, besides the outlier, for a single value of *x* there are ten different values for *y*. This pair of variables does not fit the CCC assumption: the two clusters formed with *x* (approximately separated by *x* = 13) do not match the three clusters formed with *y*. The Pearson’s correlation coefficient is the same across all these Anscombe’s examples (*p* = 0.82), whereas Spearman is 0.50 or greater. These simulated datasets show that both Pearson and Spearman are powerful in detecting linear patterns. However, any deviation in this assumption (like nonlinear relationships or outliers) affects their robustness.

We simulated additional types of relationships (Figure 1, second row), including some previously described from gene expression data [37,38,39]. For the random/independent pair of variables, all coefficients correctly agree with a value close to zero. The non-coexistence pattern, captured by all coefficients, represents a case where one gene (*x*) might be expressed while the other one (*y*) is inhibited, highlighting a potentially strong biological relationship (such as a microRNA negatively regulating another gene). For the other two examples (quadratic and two-lines), Pearson and Spearman do not capture the nonlinear pattern between variables *x* and *y*. These patterns also show how CCC uses different degrees of complexity to capture the relationships. For the quadratic pattern, for example, CCC separates *x* into more clusters (four in this case) to reach the maximum ARI. The two-lines example shows two embedded linear relationships with different slopes, which neither Pearson nor Spearman detect (*p* = −0.12 and *s* = 0.05, respectively). Here, CCC increases the complexity of the model by using eight clusters for *x* and six for *y*, resulting in *c* = 0.31.

### The CCC reveals linear and nonlinear patterns in human transcriptomic data

We next examined the characteristics of these correlation coefficients in gene expression data from GTEx v8 across different tissues. We selected the top 5,000 genes with the largest variance for our initial analyses on whole blood and then computed the correlation matrix between genes using Pearson, Spearman and CCC (see <u>Methods</u>).

We examined the distribution of each coefficient’s absolute values in GTEx (Figure 2). CCC (mean=0.14, median=0.08, sd=0.15) has a much more skewed distribution than Pearson (mean=0.31, median=0.24, sd=0.24) and Spearman (mean=0.39, median=0.37, sd=0.26). The coefficients reach a cumulative set containing 70% of gene pairs at different values (Figure 2 b), *c* = 0.18, *p* = 0.44 and *s* = 0.56, suggesting that for this type of data, the coefficients are not directly comparable by magnitude, so we used ranks for further comparisons. In GTEx v8, CCC values were closer to Spearman and vice versa than either was to Pearson (Figure 2 c). We also compared the Maximal Information Coefficient (MIC) in this data (see <u>Supplementary Note 1</u>). We found that CCC behaved very similarly to MIC, although CCC was up to two orders of magnitude faster to run (see <u>Supplementary Note 2</u>). MIC, an advanced correlation coefficient able to capture general patterns beyond linear relationships, represented a significant step forward in correlation analysis research and has been successfully used in various application domains [4,26,27]. These results suggest that our findings for CCC generalize to MIC, therefore, in the subsequent analyses we focus on CCC and linear-only coefficients.

**Figure 2:**
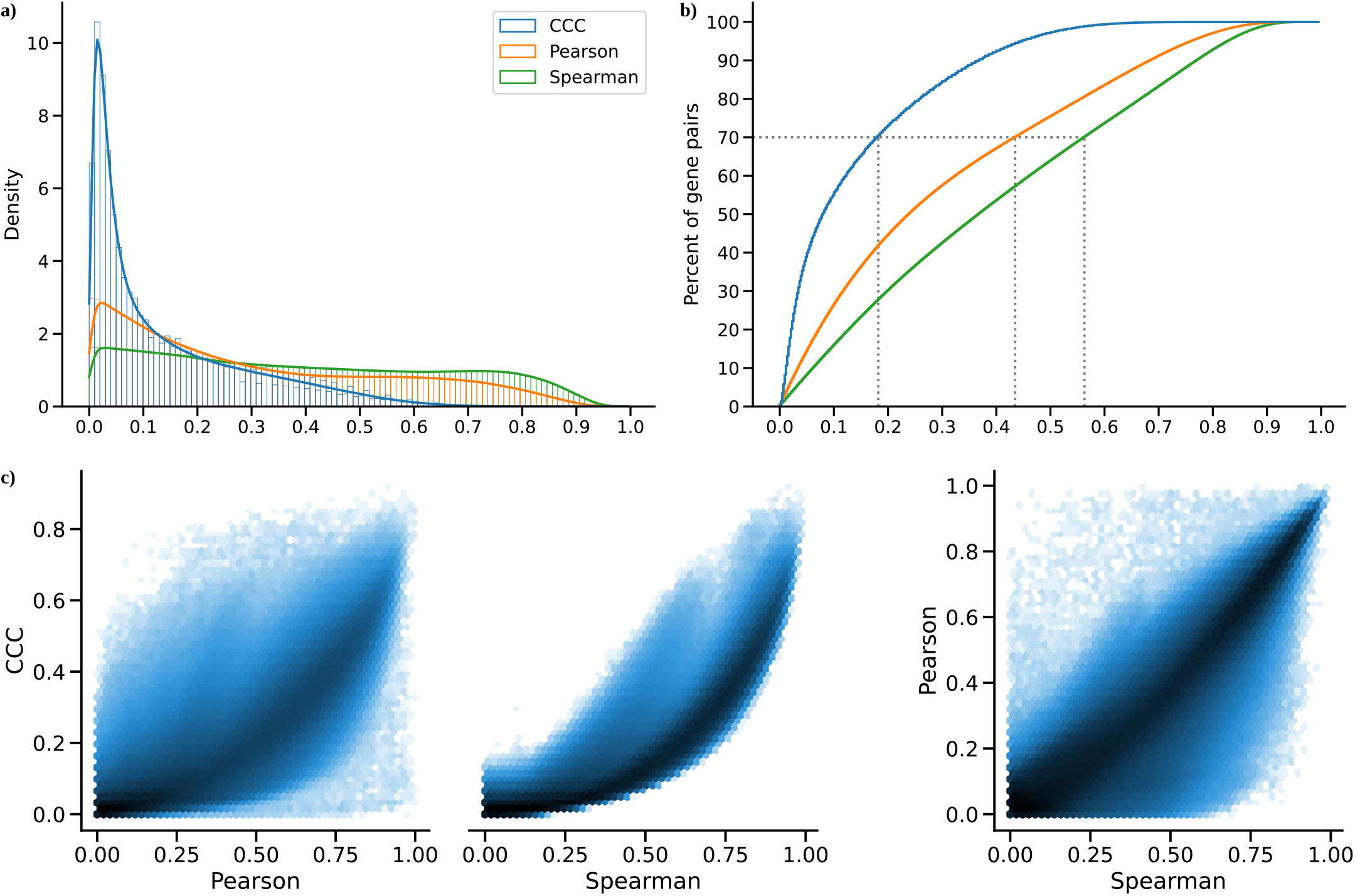
Distribution of coefficient values on gene expression (GTEx v8, whole blood). **a)** Histogram of coefficient values. **b)** Corresponding cumulative histogram. The dotted line maps the coefficient value that accumulates 70% of gene pairs. **c)** 2D histogram plot with hexagonal bins between all coefficients, where a logarithmic scale was used to color each hexagon.

A closer inspection of gene pairs that were either prioritized or disregarded by these coefficients revealed that they captured different patterns. We analyzed the agreements and disagreements by obtaining, for each coefficient, the top 30% of gene pairs with the largest correlation values (“high” set) and the bottom 30% (“low” set), resulting in six potentially overlapping categories. For most cases (76.4%), an UpSet analysis [40] (Figure 3 a) showed that the three coefficients agreed on whether there is a strong correlation (42.1%) or there is no relationship (34.3%). Since Pearson and Spearman are linear-only, and CCC can also capture these patterns, we expect that these concordant gene pairs represent clear linear patterns. CCC and Spearman agree more on either highly or poorly correlated pairs (4.0% in “high”, and 7.0% in “low”) than any of these with Pearson (all between 0.3%-3.5% for “high”, and 2.8%-5.5% for “low”). In summary, CCC agrees with either Pearson or Spearman in 90.5% of gene pairs by assigning a high or a low correlation value.

**Figure 3:**
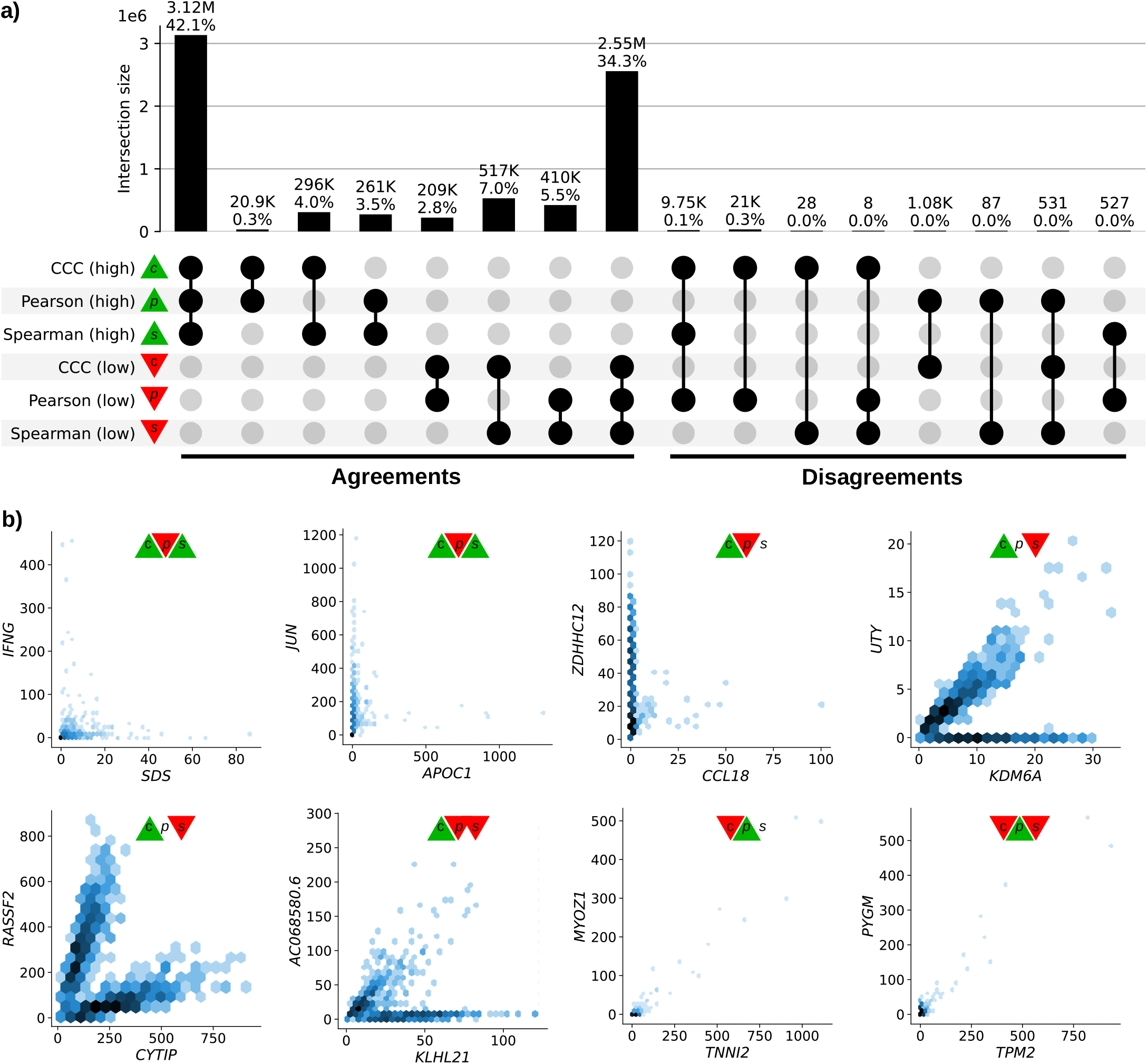
Intersection of gene pairs with high and low correlation coefficient values (GTEx v8, whole blood). **a)** UpSet plot with six categories (rows) grouping the 30% of the highest (green triangle) and lowest (red triangle) values for each coefficient. Columns show different intersections of categories grouped by agreements and disagreements. **b)** Hexagonal binning plots with examples of gene pairs where CCC (*c*) disagrees with Pearson (*p*) and Spearman (*s*). For each method, colors in the triangles indicate if the gene pair is among the top (green) or bottom (red) 30% of coefficient values. No triangle means that the correlation value for the gene pair is between the 30th and 70th percentiles (neither low nor high). A logarithmic scale was used to color each hexagon.

While there was broad agreement, more than 20,000 gene pairs with a high CCC value were not highly ranked by the other coefficients (right part of Figure 3 a). There were also gene pairs with a high Pearson value and either low CCC (1,075), low Spearman (87) or both low CCC and low Spearman values (531). However, our examination suggests that many of these cases appear to be driven by potential outliers (Figure 3 b, and analyzed later). We analyzed gene pairs among the top five of each intersection in the “Disagreements” group (Figure 3 a, right) where CCC disagrees with Pearson, Spearman or both.

The first three gene pairs at the top (*IFNG* - *SDS, JUN* - *APOC1*, and *ZDHHC12* - *CCL18*), with high CCC and low Pearson values, appear to follow a non-coexistence relationship: in samples where one of the genes is highly (slightly) expressed, the other is slightly (highly) activated, suggesting a potentially inhibiting effect. The following three gene pairs (*UTY* - *KDM6A, RASSF2* - *CYTIP*, and *AC068580*.*6* - *KLHL21*) follow patterns combining either two linear or one linear and one independent relationships. In particular, genes *UTY* and *KDM6A* (paralogs) show a nonlinear relationship where a subset of samples follows a robust linear pattern and another subset has a constant (independent) expression of one gene. This relationship is explained by the fact that *UTY* is in chromosome Y (Yq11) whereas *KDM6A* is in chromosome X (Xp11), and samples with a linear pattern are males, whereas those with no expression for *UTY* are females. This combination of linear and independent patterns is captured by CCC (*c* = 0.29, above the 80th percentile) but not by Pearson (*p* = 0.24, below the 55th percentile) or Spearman (*s* = 0.10, below the 15th percentile). Furthermore, the same gene pair pattern is highly ranked by CCC in all other tissues in GTEx, except for female-specific organs (Figure 4).

**Figure 4:**
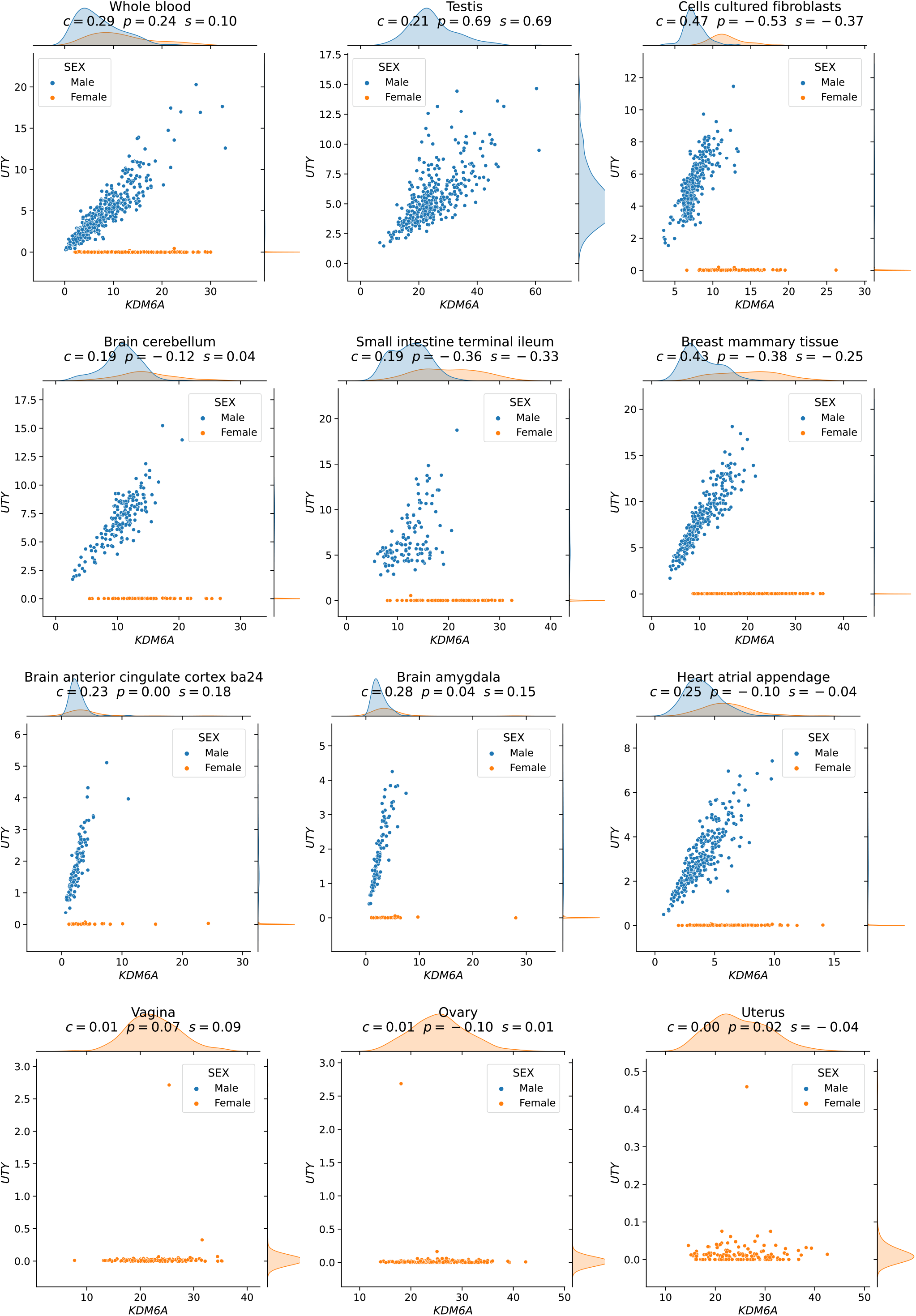
The expression levels of *KDM6A* and *UTY* display sex-specific associations across GTEx tissues. CCC captures this nonlinear relationship in all GTEx tissues (nine examples are shown in the first three rows), except in female-specific organs (last row).

### Replication of gene associations using tissue-specific gene networks from GIANT

We sought to systematically analyze discrepant scores to assess whether associations were replicated in other datasets besides GTEx. This is challenging and prone to bias because linear-only correlation coefficients are usually used in gene co-expression analyses. We used 144 tissue-specific gene networks from the Genome-wide Analysis of gene Networks in Tissues (GIANT) [41,42], where nodes represent genes and each edge a functional relationship weighted with a probability of interaction between two genes (see <u>Methods</u>). Importantly, the version of GIANT used in this study did not include GTEx samples [43], making it an ideal case for replication. These networks were built from expression and different interaction measurements, including protein-interaction, transcription factor regulation, chemical/genetic perturbations and microRNA target profiles from the Molecular Signatures Database (MSigDB [44]). We reasoned that highly-ranked gene pairs using three different coefficients in a single tissue (whole blood in GTEx, Figure 3) that represented real patterns should often replicate in a corresponding tissue or related cell lineage using the multi-cell type functional interaction networks in GIANT. In addition to predicting a network with interactions for a pair of genes, the GIANT web application can also automatically detect a relevant tissue or cell type where genes are predicted to be specifically expressed (the approach uses a machine learning method introduced in [45] and described in Methods). For example, we obtained the networks in blood and the automatically-predicted cell type for gene pairs *RASSF2* - *CYTIP* (CCC high, Figure 5 a) and *MYOZ1* - *TNNI2* (Pearson high, Figure 5 b). In addition to the gene pair, the networks include other genes connected according to their probability of interaction (up to 15 additional genes are shown), which allows estimating whether genes are part of the same tissue-specific biological process. Two large black nodes in each network’s top-left and bottom-right corners represent our gene pairs. A green edge means a close-to-zero probability of interaction, whereas a red edge represents a strong predicted relationship between the two genes. In this example, genes *RASSF2* and *CYTIP* (Figure 5 a), with a high CCC value (*c* = 0.20, above the 73th percentile) and low Pearson and Spearman (*p* = 0.16 and *s* = 0.11, below the 38th and 17th percentiles, respectively), were both strongly connected to the blood network, with interaction scores of at least 0.63 and an average of 0.75 and 0.84, respectively (Supplementary Table 1). The autodetected cell type for this pair was leukocytes, and interaction scores were similar to the blood network (Supplementary Table 1). However, genes *MYOZ1* and *TNNI2*, with a very high Pearson value (*p* = 0.97), moderate Spearman (*s* = 0.28) and very low CCC (*c* = 0.03), were predicted to belong to much less cohesive networks (Figure 5 b), with average interaction scores of 0.17 and 0.22 with the rest of the genes, respectively. Additionally, the autodetected cell type (skeletal muscle) is not related to blood or one of its cell lineages. These preliminary results suggested that CCC might be capturing blood-specific patterns missed by the other coefficients.

**Figure 5:**
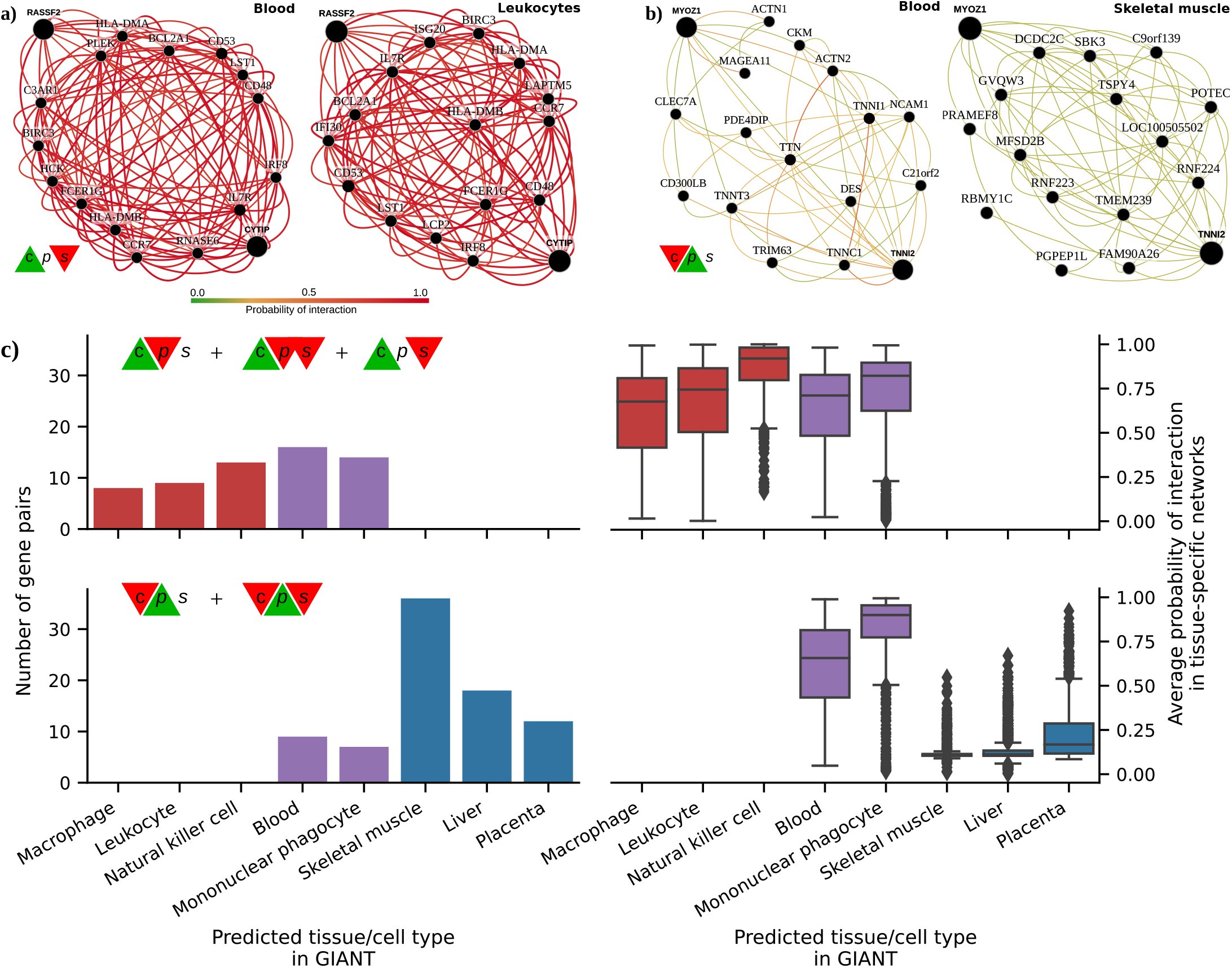
Analysis of GIANT tissue-specific predicted networks for gene pairs prioritized by correlation coefficients. **a-b)** Two gene pairs prioritized by correlation coefficients (from Figure 3 b) with their predicted networks in blood (left) and an automatically selected tissue/cell type (right) using the method described in [45]. A node represents a gene and an edge the probability that two genes are part of the same biological process in a specific cell type. A maximum of 15 genes are shown for each network. The GIANT web application automatically determined a minimum interaction confidence (edges’ weights) to be shown. These networks can be analyzed online using the following links: *RASSF2* - *CYTIP* [46], *MYOZ1* - *TNNI2* [47]. **c)** Summary of predicted tissue/cell type networks for gene pairs exclusively prioritized by CCC and Pearson. The first row combines all gene pairs where CCC is high and Pearson or Spearman are low. The second row combines all gene pairs where Pearson is high and CCC or Spearman are low. Bar plots (left) show the number of gene pairs for each predicted tissue/cell type. Box plots (right) show the average probability of interaction between genes in these predicted tissue-specific networks. Red indicates CCC-only tissues/cell types, blue are Pearson-only, and purple are shared.

We next performed a systematic evaluation using the top 100 discrepant gene pairs between CCC and the other two coefficients. For each gene pair prioritized in GTEx (whole blood), we autodetected a relevant cell type using GIANT to assess whether genes were predicted to be specifically expressed in a blood-relevant cell lineage. For this, we used the top five most commonly autodetected cell types for each coefficient and assessed connectivity in the resulting networks (see <u>Methods</u>). The top 5 predicted cell types for gene pairs highly ranked by CCC and not by the rest were all blood-specific (Figure 5 c, top left), including macrophage, leukocyte, natural killer cell, blood and mononuclear phagocyte. The average probability of interaction between genes in these CCC-ranked networks was significantly higher than the other coefficients (Figure 5 c, top right), with all medians larger than 67% and first quartiles above 41% across predicted cell types. In contrast, most Pearson’s gene pairs were predicted to be specific to tissues unrelated to blood (Figure 5 c, bottom left), with skeletal muscle being the most commonly predicted tissue. The interaction probabilities in these Pearson-ranked networks were also generally lower than in CCC, except for blood-specific gene pairs (Figure 5 c, bottom right). The associations exclusively detected by CCC in whole blood from GTEx were more strongly replicated in these independent networks that incorporated multiple data modalities. CCC-ranked gene pairs not only had high probabilities of belonging to the same biological process but were also predicted to be specifically expressed in blood cell lineages. Conversely, most Pearson-ranked gene pairs were not predicted to be blood-specific, and their interaction probabilities were relatively low. This lack of replication in GIANT suggests that top Pearson-ranked gene pairs in GTEx might be driven mainly by outliers, which is consistent with our earlier observations of outlier-driven associations (Figure 3 b).

## Discussion

We introduce the Clustermatch Correlation Coefficient (CCC), an efficient not-only-linear machine learning-based statistic. Applying CCC to GTEx v8 revealed that it was robust to outliers and detected linear relationships as well as complex and biologically meaningful patterns that standard coefficients missed. In particular, CCC alone detected gene pairs with complex nonlinear patterns from the sex chromosomes, highlighting the way that not-only-linear coefficients can play in capturing sex-specific differences. The ability to capture these nonlinear patterns, however, extends beyond sex differences: it provides a powerful approach to detect complex relationships where a subset of samples or conditions are explained by other factors (such as differences between health and disease). We found that top CCC-ranked gene pairs in whole blood from GTEx were replicated in independent tissue-specific networks trained from multiple data types and attributed to cell lineages from blood, even though CCC did not have access to any cell lineage-specific information. This suggests that CCC can disentangle intricate cell lineage-specific transcriptional patterns missed by linear-only coefficients. In addition to capturing nonlinear patterns, the CCC was more similar to Spearman than Pearson, highlighting their shared robustness to outliers. The CCC results were concordant with MIC, but much faster to compute and thus practical for large datasets. Another advantage over MIC is that CCC can also process categorical variables together with numerical values. CCC is conceptually easy to interpret and has a single parameter that controls the maximum complexity of the detected relationships while also balancing compute time.

Datasets such as Anscombe or “Datasaurus” highlight the value of visualization instead of relying on simple data summaries. While visual analysis is helpful, for many datasets examining each possible relationship is infeasible, and this is where more sophisticated and robust correlation coefficients are necessary. Advanced yet interpretable coefficients like CCC can focus human interpretation on patterns that are more likely to reflect real biology. The complexity of these patterns might reflect heterogeneity in samples that mask clear relationships between variables. For example, genes *UTY* - *KDM6A* (from sex chromosomes), detected by CCC, have a strong linear relationship but only in a subset of samples (males), which was not captured by linear-only coefficients. This example, in particular, highlights the importance of considering sex as a biological variable (SABV) [48] to avoid overlooking important differences between men and women, for instance, in disease manifestations [49,50]. More generally, a not-only-linear correlation coefficient like CCC could identify significant differences between variables (such as genes) that are explained by a third factor (beyond sex differences), that would be entirely missed by linear-only coefficients.

It is well-known that biomedical research is biased towards a small fraction of human genes [51,52]. Some genes highlighted in CCC-ranked pairs (Figure 3 b), such as *SDS* (12q24) and *ZDHHC12* (9q34), were previously found to be the focus of fewer than expected publications [53]. It is possible that the widespread use of linear coefficients may bias researchers away from genes with complex coexpression patterns. A beyond-linear gene co-expression analysis on large compendia might shed light on the function of understudied genes. For example, gene *KLHL21* (1p36) and *AC068580*.*6* (*ENSG00000235027*, in 11p15) have a high CCC value and are missed by the other coefficients. *KLHL21* was suggested as a potential therapeutic target for hepatocellular carcinoma [54] and other cancers [55,56]. Its nonlinear correlation with *AC068580*.*6* might unveil other important players in cancer initiation or progression, potentially in subsets of samples with specific characteristics (as suggested in Figure 3 b).

Not-only-linear correlation coefficients might also be helpful in the field of genetic studies. In this context, genome-wide association studies (GWAS) have been successful in understanding the molecular basis of common diseases by estimating the association between genotype and phenotype [57]. However, the estimated effect sizes of genes identified with GWAS are generally modest, and they explain only a fraction of the phenotype variance, hampering the clinical translation of these findings [58]. Recent theories, like the omnigenic model for complex traits [16,17], argue that these observations are explained by highly-interconnected gene regulatory networks, with some core genes having a more direct effect on the phenotype than others. Using this omnigenic perspective, we and others [19,20,23] have shown that integrating gene co-expression networks in genetic studies could potentially identify core genes that are missed by linear-only models alone like GWAS. Our results suggest that building these networks with more advanced and efficient correlation coefficients could better estimate gene co-expression profiles and thus more accurately identify these core genes. Approaches like CCC could play a significant role in the precision medicine field by providing the computational tools to focus on more promising genes representing potentially better candidate drug targets.

Our analyses have some limitations. We worked on a sample with the top variable genes to keep computation time feasible. Although CCC is much faster than MIC, Pearson and Spearman are still the most computationally efficient since they only rely on simple data statistics. Our results, however, reveal the advantages of using more advanced coefficients like CCC for detecting and studying more intricate molecular mechanisms that replicated in independent datasets. The application of CCC on larger compendia, such as recount3 [11] with thousands of heterogeneous samples across different conditions, can reveal other potentially meaningful gene interactions. The single parameter of CCC, *k*max, controls the maximum complexity of patterns found and also impacts the compute time. Our analysis suggested that *k*max = 10 was sufficient to identify both linear and more complex patterns in gene expression. A more comprehensive analysis of optimal values for this parameter could provide insights to adjust it for different applications or data types.

While linear and rank-based correlation coefficients are exceptionally fast to calculate, not all relevant patterns in biological datasets are linear. For example, patterns associated with sex as a biological variable are not apparent to the linear-only coefficients that we evaluated but are revealed by not-only-linear methods. Beyond sex differences, being able to use a method that inherently identifies patterns driven by other factors is likely to be desirable. Not-only-linear coefficients can also disentangle intricate yet relevant patterns from expression data alone that were replicated in models integrating different data modalities. CCC, in particular, is highly parallelizable, and we anticipate efficient GPU-based implementations that could make it even faster. The CCC is an efficient, next-generation correlation coefficient that is highly effective in transcriptome analyses and potentially useful in a broad range of other domains.

## Methods

The code needed to reproduce all of our analyses and generate the figures is available in https://github.com/greenelab/ccc. We provide scripts to download the required data and run all the steps. A Docker image is provided to use the same runtime environment.

### The CCC algorithm

The Clustermatch Correlation Coefficient (CCC) computes a similarity value *c* ∈ [0, 1] between any pair of numerical or categorical features/variables **x** and **y** measured on *n* objects. CCC assumes that if two features **x** and **y** are similar, then the partitioning by clustering of the *n* objects using each feature separately should match. For example, given **x** = (11, 27, 32, 40) and **y** = 10*x* = (110, 270, 320, 400), where *n* = 4, partitioning each variable into two clusters (*k* = 2) using their medians (29.5 for **x** and 295 for **y**) would result in partition 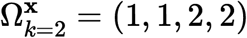 for **x**, and partition 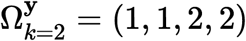 for **y**. Then, the agreement between 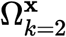 and 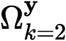 can be computed using any measure of similarity between partitions, like the adjusted Rand index (ARI) [32]. In that case, it will return the maximum value (1.0 in the case of ARI). Note that the same value of *k* might not be the right one to find a relationship between any two features. For instance, in the quadratic example in Figure 1, CCC returns a value of 0.36 (grouping objects in four clusters using one feature and two using the other). If we used only two clusters instead, CCC would return a similarity value of 0.02. Therefore, the CCC algorithm (shown below) searches for this optimal number of clusters given a maximum ***k***, which is its single parameter ***k*max**.

#### Algorithm 1: CCC algorithm

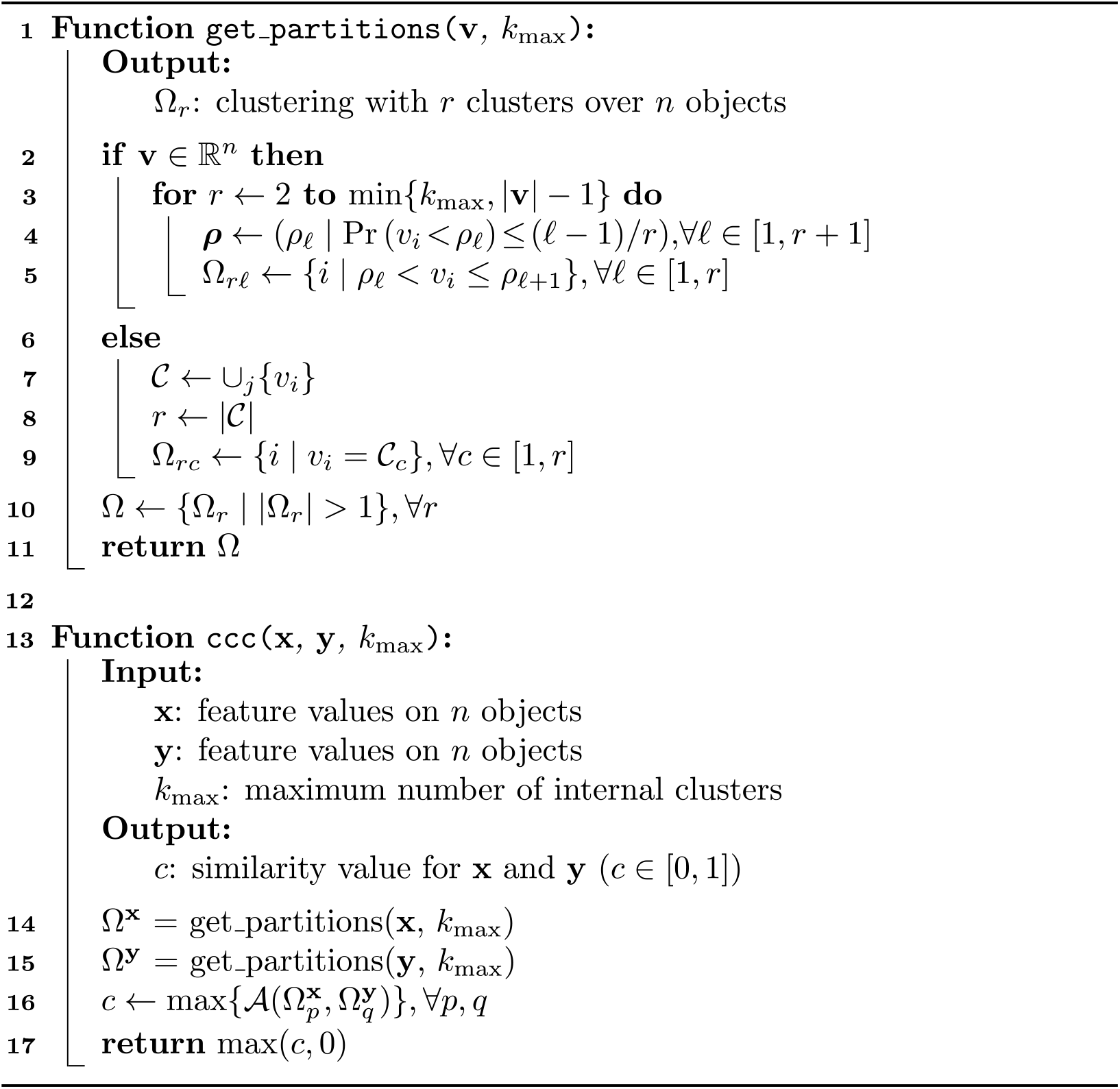

The main function of the algorithm, `ccc`, generates a list of partitionings **Ω**^**x**^ and **Ω**^**y**^ (lines 14 and 15), for each feature **x** and **y**. Then, it computes the ARI between each partition in **Ω**^**x**^ and **Ω**^**y**^ (line 16), and then it keeps the pair that generates the maximum ARI. Finally, since ARI does not have a lower bound (it could return negative values, which in our case are not meaningful), CCC returns only values between 0 and 1 (line 17).

Interestingly, since CCC only needs a pair of partitions to compute a similarity value, any type of feature that can be used to perform clustering/grouping is supported. If the feature is numerical (lines 2 to 5 in the `get_partitions` function), then quantiles are used for clustering (for example, the median generates ***k*** = 2 clusters of objects), from ***k* = 2** to k = *k*_**max**_. If the feature is categorical (lines 7 to 9), the categories are used to group objects together. Consequently, since features are internally categorized into clusters, numerical and categorical variables can be naturally integrated since clusters do not need an order.

For all our analyses we used ***k***_**max**_ = 10. This means that for each gene pair, 18 partitions are generated (9 for each gene, from ***k*** = 2 to ***k*** = 10), and 81 ARI comparisons are performed. Smaller values of ***k***_**max**_ can reduce computation time, although at the expense of missing more complex/general relationships. Our examples in Figure 1 suggest that using ***k***_**max**_ = **2** would force CCC to find linear-only patterns, which could be a valid use case scenario where only this kind of relationships are desired. In addition, ***k***_**max**_ = **2** implies that only two partitions are generated, and only one ARI comparison is performed. In this regard, our Python implementation of CCC provides flexibility in specifying ***k***_**max**_. For instance, instead of the maximum *k* (an integer), the parameter could be a custom list of integers: for example, [2, 5, 10] will partition the data into two, five and ten clusters.

For a single pair of features (genes in our study), generating partitions or computing their similarity can be parallelized. We used three CPU cores in our analyses to speed up the computation of CCC. A future improved implementation of CCC could potentially use graphical processing units (GPU) to parallelize its computation further.

A Python implementation of CCC (optimized with `numba` [59]) can be found in our Github repository [60], as well as a package published in the Python Package Index (PyPI) that can be easily installed.

### Gene expression data and preprocessing

We downloaded GTEx v8 data for all tissues, normalized using TPM (transcripts per million), and focused our primary analysis on whole blood, which has a good sample size (755). We selected the top 5,000 genes from whole blood with the largest variance after standardizing with *log*(*x* + 1) to avoid a bias towards highly-expressed genes. We then computed Pearson, Spearman, MIC and CCC on these 5,000 genes across all 755 samples on the TPM-normalized data, generating a pairwise similarity matrix of size 5,000 × 5,000.

### Tissue-specific network analyses using GIANT

We accessed tissue-specific gene networks of GIANT using both the web interface and web services provided by HumanBase [42]. The GIANT version used in this study included 987 genome-scale datasets with approximately 38,000 conditions from around 14,000 publications. Details on how these networks were built are described in [31]. Briefly, tissue-specific gene networks were built using gene expression data (without GTEx samples [43]) from the NCBI’s Gene Expression Omnibus (GEO) [61], protein-protein interaction (BioGRID [62], IntAct [63], MINT [64] and MIPS [65]), transcription factor regulation using binding motifs from JASPAR [66], and chemical and genetic perturbations from MSigDB [67]. Gene expression data were log-transformed, and the Pearson correlation was computed for each gene pair, normalized using the Fisher’s z transform, and z-scores discretized into different bins. Gold standards for tissue-specific functional relationships were built using expert curation and experimentally derived gene annotations from the Gene Ontology. Then, one naive Bayesian classifier (using C++ implementations from the Sleipnir library [68]) for each of the 144 tissues was trained using these gold standards. Finally, these classifiers were used to estimate the probability of tissue-specific interactions for each gene pair.

For each pair of genes prioritized in our study using GTEx, we used GIANT through HumanBase to obtain 1) a predicted gene network for blood (manually selected to match whole blood in GTEx) and 2) a gene network with an automatically predicted tissue using the method described in [45] and provided by HumanBase web interfaces/services. Briefly, the tissue prediction approach trains a machine learning model using comprehensive transcriptional data with human-curated markers of different cell lineages (e.g., macrophages) as gold standards. Then, these models are used to predict other cell lineage-specific genes. In addition to reporting this predicted tissue or cell lineage, we computed the average probability of interaction between all genes in the network retrieved from GIANT. Following the default procedure used in GIANT, we included the top 15 genes with the highest probability of interaction with the queried gene pair for each network.

### Maximal Information Coefficient (MIC)

We used the Python package `minepy` [69,70] (version 1.2.5) to estimate the MIC coeicient. In GTEx v8 (whole blood), we used MIC_e_ (an improved implementation of the original MIC introduced in [71]) with the default parameters `alpha=0.6`, `c=15` and `estimator=‘mic_e’`. We used the `pairwise_distances` function from `scikit-learn` [72] to parallelize the computation of MIC on GTEx. For our computational complexity analyses (see <u>Supplementary Material</u>), we ran the original MIC (using parameter `estimator=‘mic_approx’`) and MIC_e_ (`estimator=‘mic_e’`).

## Supplementary material

### Supplementary Note 1: Comparison with the Maximal Information Coefficient (MIC) on gene expression data

We compared all the coefficients in this study with MIC [24], a popular nonlinear method that can find complex relationships in data, although very computationally intensive [73]. We ran MIC_e_ (see Methods) on all possible pairwise comparisons of our 5,000 highly variable genes from whole blood in GTEx v8. This took 4 days and 19 hours to finish (compared with 9 hours for CCC). Then we performed the analysis on the distribution of coefficients (the same as in the main text), shown in Figure 6. We verified that CCC and MIC behave similarly in this dataset, with essentially the same distribution but only shifted. Figure 6 c shows that these two coefficients relate almost linearly, and both compare very similarly with Pearson and Spearman.

**Figure 6:**
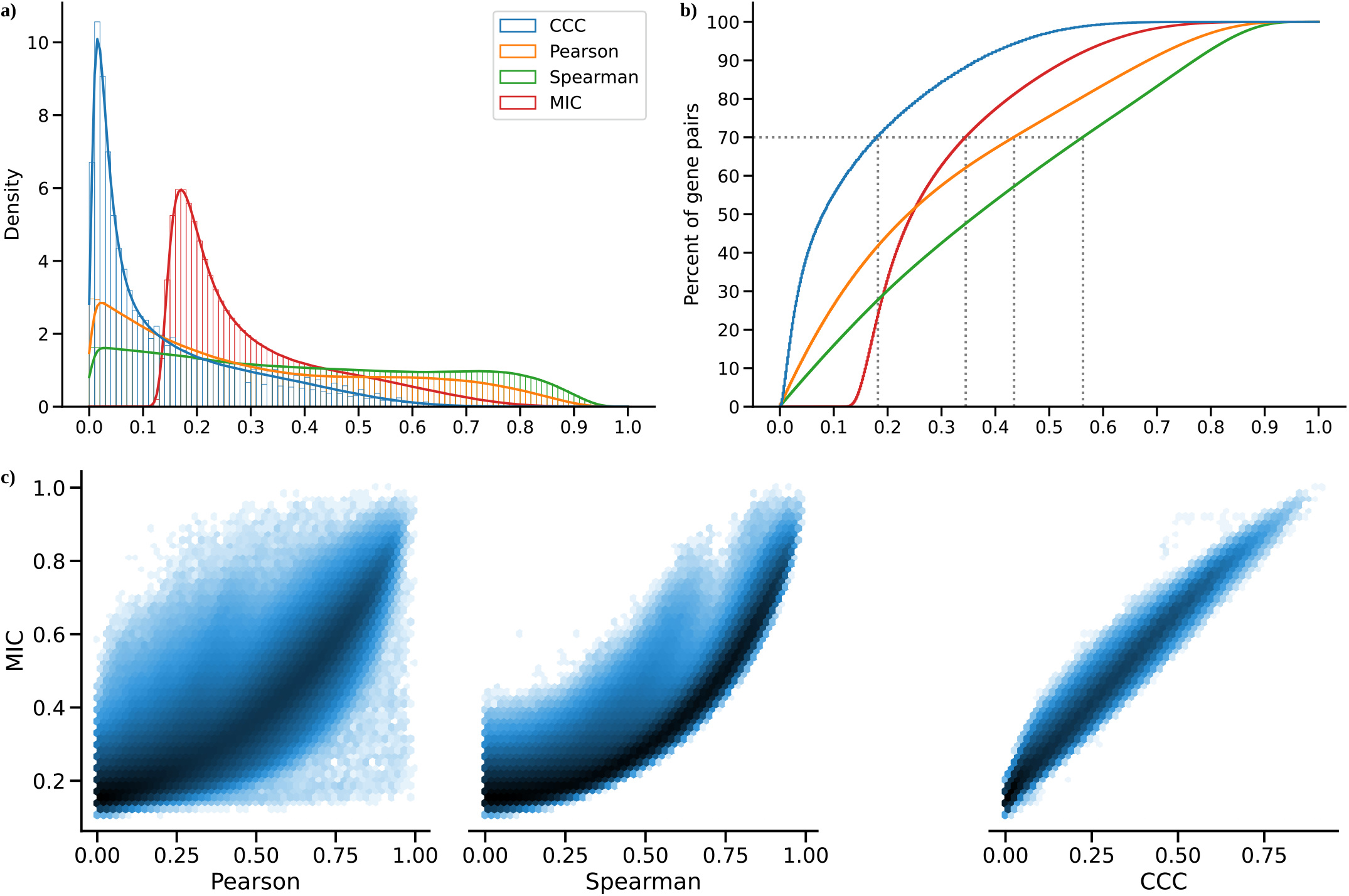
Distribution of MIC values on gene expression (GTEx v8, whole blood) and comparison with other methods. **a)** Histogram of coefficient values. **b)** Corresponding cumulative histogram. The dotted line maps the coefficient value that accumulates 70% of gene pairs. **c)** 2D histogram plot with hexagonal bins between all coefficients, where a logarithmic scale was used to color each hexagon.

### Supplementary Note 2: Computational complexity of coefficients

We also compared CCC with the other coefficients in terms of computational complexity. Although CCC and MIC might identify similar gene pairs in gene expression data (see <u>here</u>), the use of MIC in large datasets remains limited due to its very long computation time, despite some methodological/implementation improvements [69,73,74,75,76]. The original MIC implementation uses ApproxMaxMI, a computationally demanding heuristic estimator [37]. Recently, a more efficient implementation called MIC_e_ was proposed [71]. These two MIC estimators are provided by the `minepy` package [69], a C implementation available for Python. We compared all these coefficients in terms of computation time on randomly generated variables of different sizes, which simulates a scenario of gene expression data with different numbers of conditions. Differently from the rest, CCC allows us to easily parallelize the computation of a single gene pair (see <u>Methods</u>), so we also tested the cases using 1 and 3 CPU cores. Figure 7 shows the time in seconds in log scale.

**Figure 7:**
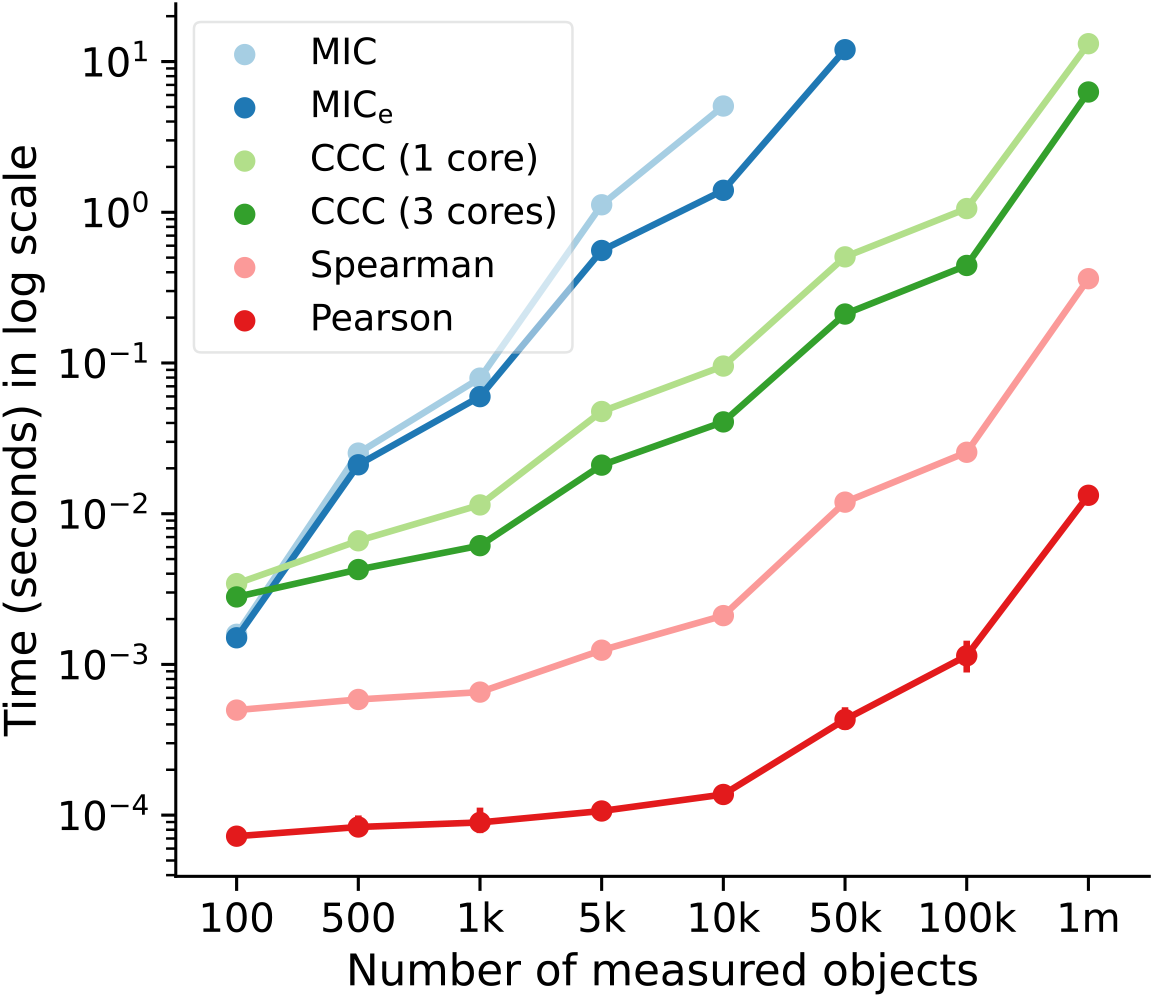
Computational complexity of all correlation coefficients on simulated data. We simulated variables/features with varying data sizes (from 100 to a million, *x*-axis). The plot shows the average time in seconds (log-scale) taken for each coefficient on ten repetitions (1000 repetitions were performed for data size 100). CCC was run using 1 and 3 CPU cores. MIC and MIC_e_ did not finish running in a reasonable amount of time for data sizes of 10,000 and 100,000, respectively.

As we already expected, Pearson and Spearman were the fastest, given that they only need to compute basic summary statistics from the data. For example, Pearson is three orders of magnitude faster than CCC. Among the nonlinear coefficients, CCC was faster than the two MIC variations (up to two orders of magnitude), with the only exception in very small data sizes. The difference is important because both MIC variants were implemented in C [69], a high-performance programming language, whereas CCC was implemented in Python (optimized with `numba`). For a data size of a million, the multi-core CCC was twice as fast as the single-core CCC. This suggests that new implementations using more advanced processing units (such as GPUs) are feasible and could make CCC reach speeds closer to Pearson.

## Tissue-specific gene networks with GIANT

**Table 1:**
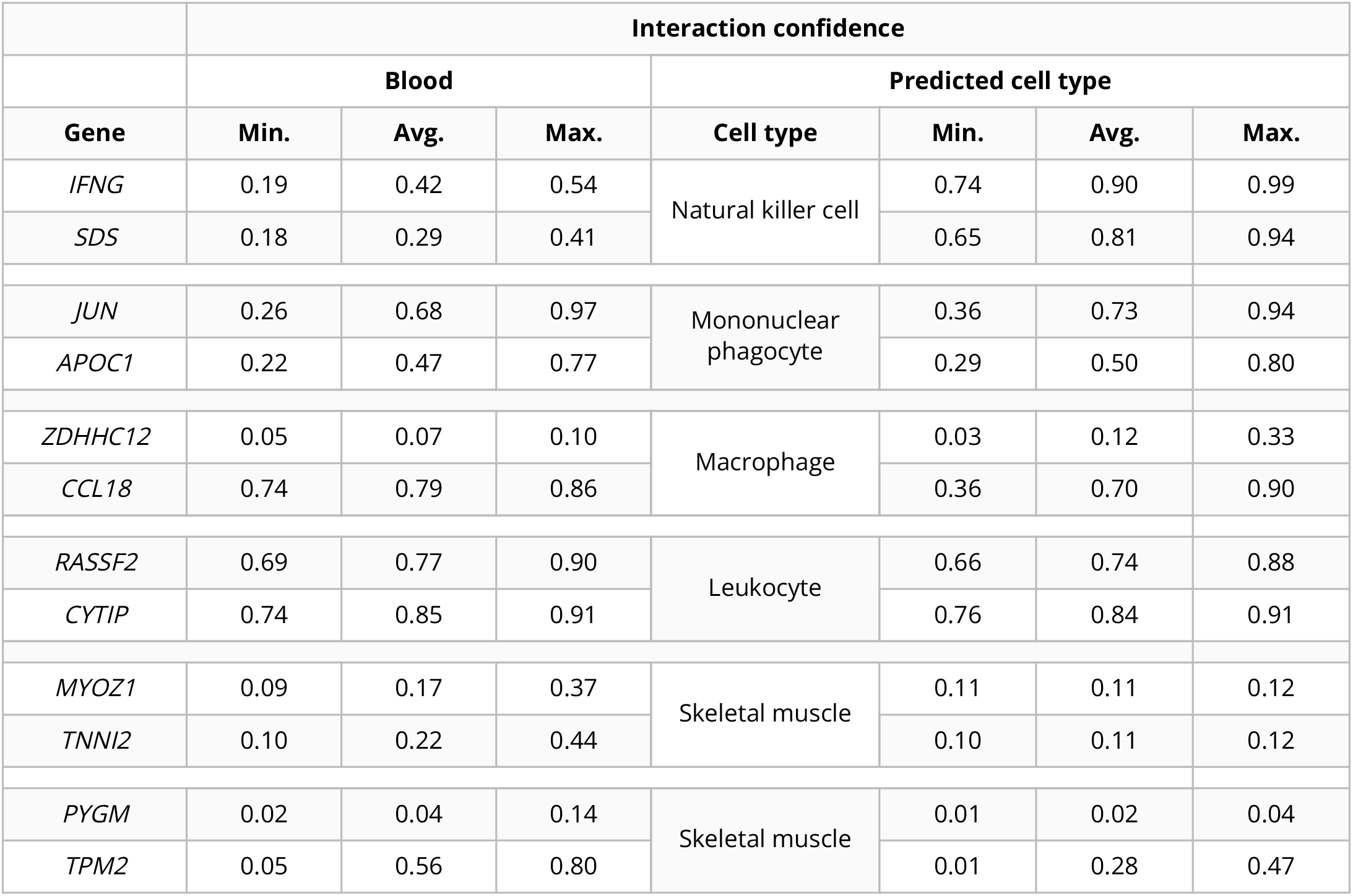
Network statistics of six gene pairs shown in Figure 3 b for blood and predicted cell types. Only gene pairs present in GIANT models are listed. For each gene in the pair (first column), the minimum, average and maximum interaction coefficients with the other genes in the network are shown.

|  | Interaction confidence |  |  |  |  |  |  |
| --- | --- | --- | --- | --- | --- | --- | --- |
|  | Blood |  |  | Predicted cell type |  |  |  |
| Gene | Min. | Avg. | Max. | Cell type | Min. | Avg. | Max. |
| *IFNG* | 0.19 | 0.42 | 0.54 | Natural killer cell | 0.74 | 0.90 | 0.99 |
| *SDS* | 0.18 | 0.29 | 0.41 |  | 0.65 | 0.81 | 0.94 |
| *JUN* | 0.26 | 0.68 | 0.97 | Mononuclear phagocyte | 0.36 | 0.73 | 0.94 |
| *APOC1* | 0.22 | 0.47 | 0.77 |  | 0.29 | 0.50 | 0.80 |
| *ZDHHC12* | 0.05 | 0.07 | 0.10 | Macrophage | 0.03 | 0.12 | 0.33 |
| *CCL18* | 0.74 | 0.79 | 0.86 |  | 0.36 | 0.70 | 0.90 |
| *RASSF2* | 0.69 | 0.77 | 0.90 | Leukocyte | 0.66 | 0.74 | 0.88 |
| *CYTIP* | 0.74 | 0.85 | 0.91 |  | 0.76 | 0.84 | 0.91 |
| *MYOZ1* | 0.09 | 0.17 | 0.37 | Skeletal muscle | 0.11 | 0.11 | 0.12 |
| *TNNI2* | 0.10 | 0.22 | 0.44 |  | 0.10 | 0.11 | 0.12 |
| *PYGM* | 0.02 | 0.04 | 0.14 | Skeletal muscle | 0.01 | 0.02 | 0.04 |
| *TPM2* | 0.05 | 0.56 | 0.80 |  | 0.01 | 0.28 | 0.47 |

## Notes

https://github.com/greenelab/ccc

https://github.com/greenelab/ccc-manuscript

### Competing Interest Statement

The authors have declared no competing interest.

